# Complex community-wide consequences of consumer sexual dimorphism

**DOI:** 10.1101/634782

**Authors:** Stephen P. De Lisle, Sebastian J. Schrieber, Daniel I. Bolnick

**Affiliations:** Department of Ecology & Evolutionary Biology, University of Connecticut, Storrs, CT 06269; Evolutionary Ecology Unit, Department of Biology, Lund University, Sölvegatan 37, 22362 Lund Sweden; Department of Evolution and Ecology and Center for Population Biology, University of California, Davis, CA 95616

**Keywords:** apparent competition, community assembly, competitive exclusion, ecological sexual dimorphism, resource competition

## Abstract

Sexual dimorphism is a ubiquitous source of within-species variation, yet the community-level consequences of sex differences remain poorly understood. Here, we analyze a bitrophic model of two competing resource species and a sexually-reproducing consumer species. We show that consumer sex differences in resource acquisition can have striking consequences for consumer-resource coexistence, abundance, and dynamics. Under both direct interspecific competition and apparent competition between two resource species, sexual dimorphism in consumers’ attack rates can mediate coexistence of the resource species, while in other cases can lead to exclusion when stable coexistence is typically expected. Slight sex differences in total resource acquisition also can reverse competitive outcomes and lead to density cycles. These effects are expected whenever both consumer sexes require different amounts or types of resources to reproduce. Our results suggest that consumer sexual dimorphism, which is common, has wide-reaching implications for the assembly and dynamics of natural communities.

**Statement of authorship:** DB SD and SJS designed the study, SJS performed the mathematical analysis, SD performed the simulations and drafted the manuscript. All authors revised the manuscript.

**Data accessibility statement:** No data is used

## Introduction

Within-species variation is one key feature of natural populations that has emerged as a critical contributor to community ecology and species interactions (Hughes et al. 2008, Bolnick et al. 2011, Des Roches et al. 2018). Even in the simplest scenario of a single-species community with a single optimum phenotype, within-population variation is expected to reduce population growth rate of a well-adapted population via effects on mean fitness (Haldane 1937). In more complex multi-species communities, phenotypic variation in consumers or their resources can either promote or constrain coexistence between competing species, under many circumstances yielding different conclusions than would be reached from models ignoring within species variation (Bolnick et al. 2011, Schreiber et al. 2011, Patel and Schreiber 2015, Cortez and Patel 2017). One emerging question (Bolnick et al. 2011) from this work is the degree to which understanding the specific source of phenotypic variation matters in community ecology.

Here, we explore the ecological consequences of sexual dimorphism, a central feature of metazoans and many plant populations that has not been fully incorporated into ecological theory. The gamete differences that define the sexes are expected to lead to divergence between males and females in a suite of life history traits (Trivers 1972, Shärer et al. 2012), and phenotypic traits. This divergence is iconically manifest as striking sexual dimorphism in sexually-selected traits such as body size or courtship displays (Darwin 1871, Andersson 1994). Although sexual selection and associated mate choice behaviors themselves may have relevance to interspecific ecological interactions (Gomez-Llano et al. 2018), the different life histories that define the sexes are often expected lead to different nutritional and resource requirements for males and females (Maklakov et al. 2008). As a specific example, in insects, differential contributions of longevity and fecundity to male and female lifetime reproductive success result in different combinations of macronutrients that maximize male and female fitness (Maklakov et al. 2008, Reddiex et al. 2013, Garlapow et al. 2015, Jensen et al. 2015, Camus et al. 2017). Similarly, nutritional models of optimal foraging have been proposed to explain sex differences in diets of moose (Belovksy 1978). Consequently, males and females of many species have evolved divergent resource use, either as an indirect outcome of divergent reproductive roles or through other forms of sex-specific natural selection, resulting in sex-differences in diet composition and often in corresponding trophic morphology (Slatkin 1984, Temeles 1985, Shine 1989, Temeles et al. 2000, De Lisle and Rowe 2015a, De Lisle 2019).

Data suggest that these ‘ecological’ sexual dimorphisms can have substantial consequences for community structure (Fryxell et al. 2015, Pincheira-Donoso et al. 2018, Start and De Lisle 2018, Tsuji and Fukami 2018, 2020). More generally, sex differences may play an important role in the relationship between ecological and evolutionary dynamics (Giery and Layman 2019, Svensson 2019, Fryxell et al. 2019). Sexual dimorphism thus represents a key source of ecologically-relevant variation within species. Within-species variation in resource specialization is commonplace (Bolnick et al. 2003), with important consequences for species interactions and community assembly (Hughes et al. 2008, Bolnick et al. 2011, Des Roches et al. 2018). Critically, as a source of intraspecific variation, sexual dimorphism may have different consequences from other types of variation. Unlike most phenotypes, whose relative abundances can evolve to reflect local ratios of alternative resources, the ratio of males to females (at birth) is expected to be maintained at 1:1 (Darwin 1871, Fisher 1930), though their phenotypes and demography may diverge to have differential ecological impacts. Thus, the rate at which males are born (which determines predation pressure on males’ prey), is strongly dependent on females’ foraging success. Perhaps to a lesser extent (depending on mating system), the rate of female offspring production can depend on male foraging success. Importantly, persistence of a sexually

reproducing consumer depends on the persistence of both sexes. This coupling of the dynamics of two consumer phenotypes has unknown consequences for consumer-resource dynamics, consumer mediated coexistence of the resource species, and apparent competition between the resource species. Using a simple model of consumer-resource dynamics, we show that consumer sexual dimorphism can influence (both positively and negatively) coexistence between resource species (competing directly or apparently), and species’ abundance.

## Materials and Methods

Our model represents a simple extension of a classic model of consumer-resource dynamics, in which a consumer species exploits two resources that themselves undergo density-dependent growth (Figure 1). Consumer growth rate is limited only by resource abundance. The original version of this model did not consider within-species phenotypic variation, and can lead to exclusion of one resource (‘apparent competition’; (e.g. Holt 1977). More recently, (Schreiber et al. 2011) showed that quantitative trait variation in the predator, affecting attack rates on the two resources, can facilitate coexistence between the prey. Here, we instead allow for the possibility that male and female consumers differ in resource-specific attack rates. We describe the population dynamics of two resources (with densities *R_i_, R_j_*) and consumer males (with density *M*) and females (with density *F*) using the system of ordinary differential equations:

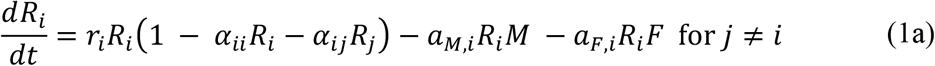

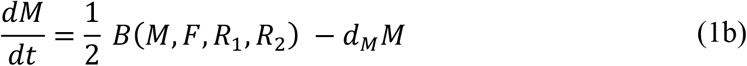

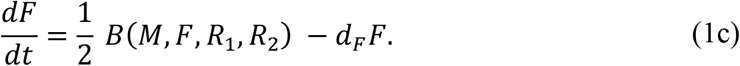

**Figure 1.**
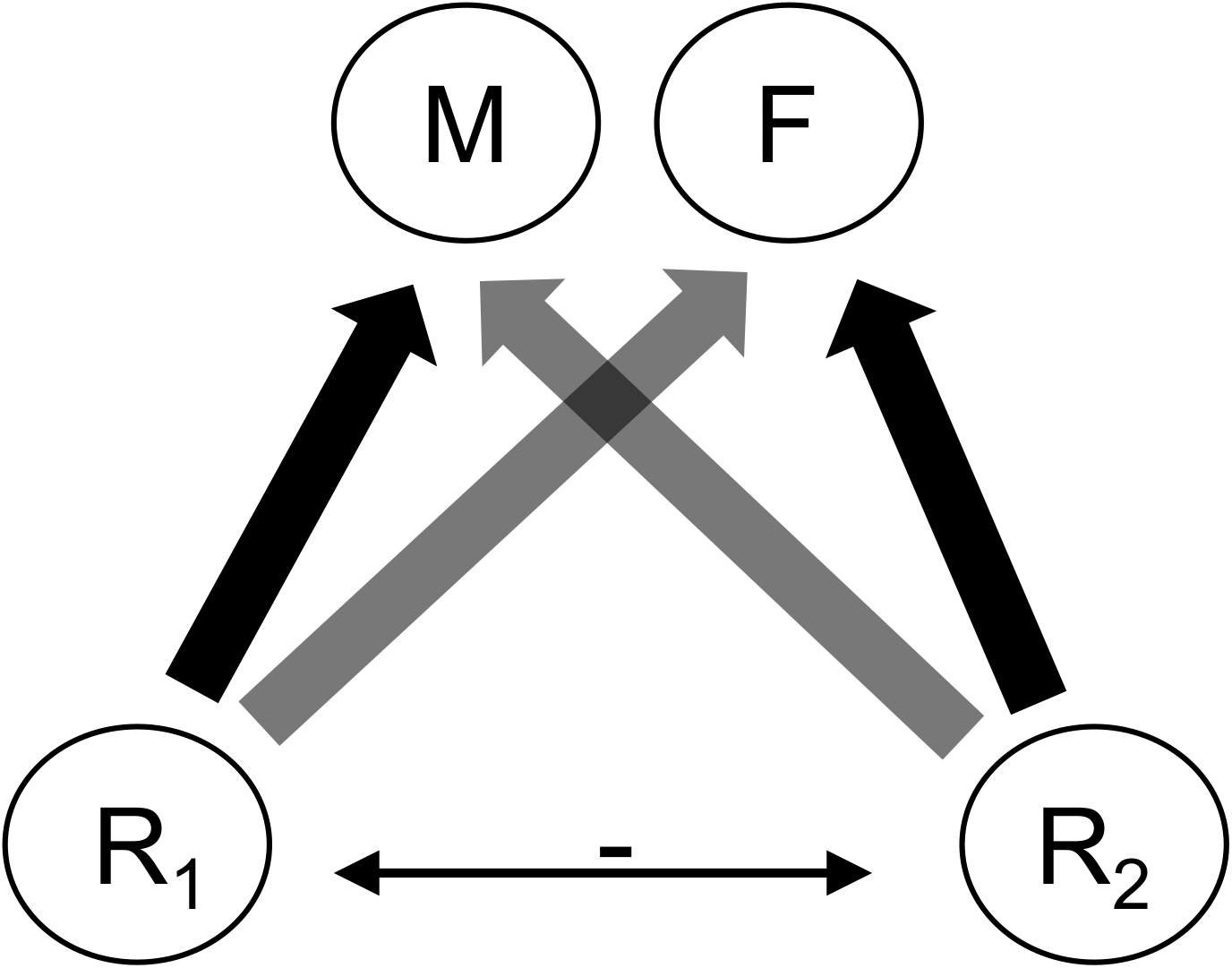
Illustration of the model structure. Males (*M*) and females (*F*) of a consumer species exploit two resources (*R*_1_ and *R*_2_) that may or may not also compete directly (double arrow). Sex-specific attack rates (black arrows) generate sex differences in ecological niche such that males and females preferentially attack *R*_1_ and *R*_2_, respectively, with varying degrees of overlap (grey arrows). Although males and females may consume different resources, fitness is equal across the sexes and so the dynamics of males, females, resource 1 and resource 2 are coupled.

Each resource undergoes density-dependent population growth with intrinsic per-capita growth rates *r,* with competition represented by competition coefficients *α*. Resources are also regulated by consumers depending on sex-specific attack rates by males *a_M_* and by females *a_F_*. Consumer dynamics are governed by females’ birth rate *B* (half of the offspring being female), and limited by sex-specific intrinsic mortality rates *d_M_* and *d_F_*.

An appropriate function describing birth rates has been a point of debate for demographers, with the general conclusion being that any function describing birth rates should capture the negative effects of extreme sex ratio skew on birth rates, with the extreme being that birth rates should be zero when one sex is absent (Caswell and Weeks 1986), or unable to breed. Generally, treating birth rates proportional to the harmonic mean density of males and females is agreed as the best approach (Caswell and Weeks 1986, Lindstöm and Kokko 1998), and is also empirically supported (Miller and Inouye 2011). Here, we extend this harmonic mean birth function to account for each sex’s foraging success, on the logic that each sex must both be present and sufficiently well-fed to reproduce:

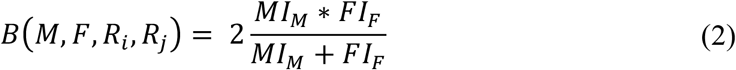

where *I* is the energy intake of a given sex:

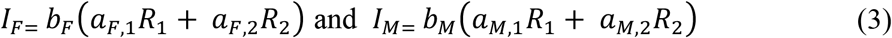

which couples the dynamics of consumers and resources. In this model the rate of consumer births depends on not just the number of males and females but also on the abundance of both prey and the prey preference of each sex, where *MI_M_* is the number of males weighted by their energy to reproduce (similarly with females), *b* is a scaling constant reflecting the degree to which sex-specific resource acquisition influences birth rate, and *a_M,i_, a_F,i_* are the attack rates on the resource *i*. Although we make no genetic assumptions, we note that our birth function, and thus our model, applies to sexually-reproducing diploid consumers. As an alternative to the inclusion of constants *b_M_* and *b_F_*, equation 2 can be equivalently expressed with the inclusion of a “harem size” parameter, *h* (Caswell and Weeks 1986, Lindstöm and Kokko 1998; see supplemental material); changes in *b_M_* relative to *b_F_* alter the degree to which each sex contributes to birth rates in an equivalent manner to the effects of deviations of *h* from unity. We focus on the parameterization in equation 2 because it illustrates that the mathematical/demographic effects of “harem size” can in fact be brought about by any factor that alters the relative contribution of sex-specific density to birth rates.

Because we are specifically interested in understanding effects of sexual dimorphism in prey preference, we relate male and female resource-specific attack rates via the degree of sexual dimorphism, *β,* such that

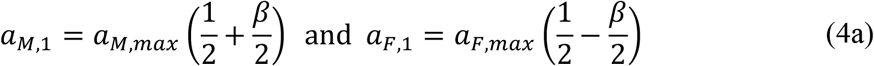

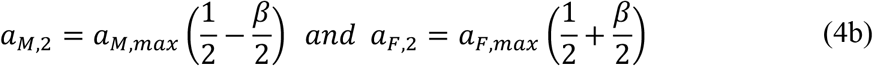

where *β* ranges from zero to one, with *β* = 0 representing complete sexual monomorphism if maximum attack rates are equal across the sexes (both sexes are generalists, attacking each prey at the rates, *a_F,max_* /2 and *a_M,max_* /2), and *β* = 1 representing complete sexual dimorphism such that males only attack *R*_1_ and females only attack *R*_2_. Sex differences in *a_max_* reflect a situation where males and females intake different total amounts of resources, independent of any difference in resource preference. In our study we explore both types of sex differences in resource acquisition; sexual dimorphism in resource preference captured by *β*, as well as sex differences in total resource acquisition captured by sex-specific *a_max_*.

By assuming that resource acquisition of males and females contributes to the consumer birth rate, we are not necessarily making any assumption about parental care or condition-dependent fecundity. Rather, this is an assumption that the fraction of each sex available for reproduction depends in part on resource acquisition, a realistic feature of many organisms. For instance, males’ energetic costs of finding a mate, defending a territory, care for offspring, or expressing a sexually-selected trait are all expected to depend on the resource pool available to males (Rowe and Houle 1996). Females’ ability to produce eggs, gestate, and provision young (e.g., lactation in mammals) similarly depends on their energy intake. Parental care and other factors can also lead to associations between sex-specific resource abundance and consumer birth rates, beyond what would be expected beyond the costs of simply being available to mate.

We analyzed this model to find equilibria when both one or two resources are present, as well as invasion criteria for a resource into a two species community at equilibrium. Using the mathematical theory of permanence (Schreiber 2000, Patel & Schreiber 2018), we can use these invasion criteria to determine whether all three species coexist (in the sense of permanence), exhibit a bistability, or competitive exclusion occurs. In addition to presenting our analytical results below, we provide further mathematical details in Supplement A. We also numerically explored the behavior of the model under a variety of scenarios using the packages deSolve v. 1.21 (Soetgart et al. 2010) and caTools v. 1.17.1.2 (Tuszynski 2019) in R v. 3.5.0 (R Core Team 2018). Importantly, using simulations allowed us to explore the behavior of our model for cases where 2-species consumer-resource equilibria do not exist. Because there are a large number of potential combinations of parameters that could be explored, we focused our simulations on three different biological scenarios: 1) completely symmetric male and female total attack rates and contributions to birth rate, 2) asymmetric total attack rates across the sexes, and 3) asymmetric contribution of male and female resource acquisition to consumer birth rates. The first scenario applies most readily to organisms with biparental care and similar total caloric requirements across the sexes. The second scenario likely applies to many organisms where total lifetime resource acquisition is higher for one sex, which is the case for many organisms where the sexes differ, for example, in body size. The third scenario is another realistic departure from the first, relevant for organisms where resource acquisition in one sex (e.g., females) has a greater influence on that sexes mating propensity, or cases with biased operational sex ratios due to polyandry or polygyny. For each scenario we explore the consequences of consumer ecological sexual dimorphism (*β*) for consumer persistence and coexistence. For each set of parameter values, we simulated the model 1,000 time steps (to ensure the simulation reaches its equilibrium).

For each run, we determined the equilibrium abundances of *M, F, R*_1_ and *R*_2_ at *t* = 1,000. We also calculated the standard deviation of species’ abundances in the last 50 time-steps. Although we present solutions simulated on the scale of *α_ii_* = 0.1, we obtained qualitatively equivalent conclusions rescaling under a wide range values of competition coefficients (see Table S1B, Supplement B). The R script to generate all figures and results presented is available in the supplementary material.

## Results

### Analytical results

Our analysis begins by considering a subsystem of a single resource species, say species i, and the consumer species. That is, we explore the conditions under which a sexually dimorphic consumer can persist on a single resource species. These two species can coexist if

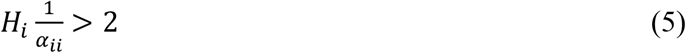

where 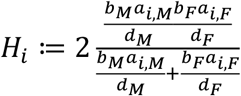 is the harmonic mean of the lifetime per-capita resource contributions of each sex to reproduction, and 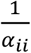 corresponds to the carrying capacity of resource species i. Intuitively, 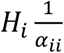 corresponds to the average number of offspring produced by a mating pair during their lifetime when the resource species is at its carrying capacity. These harmonic means decrease to zero with the degree of sexual dimorphism (i.e. *H_i_* is a decreasing function of *β* and *H_i_*=0 when *β* = 1), there always is a critical degree of sexual dimorphism above which the consumer cannot persist on a single resource species. Intuitively, this arises when one of the sexes specializes too much on the other (absent) resource species and, consequently, contributes too little to reproduction. When the coexistence condition holds, the consumer-resource species pair coexist at the following equilibrium densities:

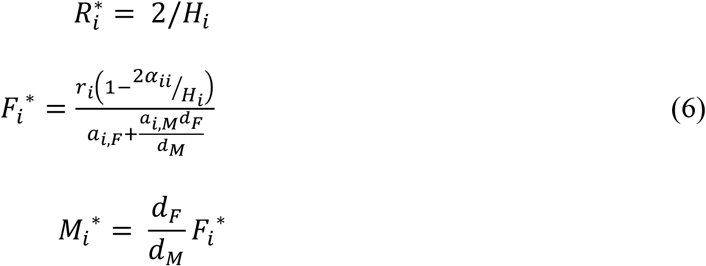

At these densities of the consumer and resource *i*, the second (rare) resource species *j* can invade if its per capita growth rate

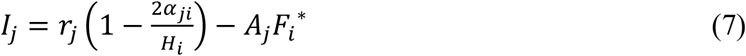

is positive. Here, *A_i_* = *a_F,j_* + *a_M,j_ d_F_*/*d_M_* is the average attack rate on resource *j* when the males and females are at an equilibrium. Provided the consumer species can persist on each of the resource species individually, the invasion growth rates *I_1_* and *I_2_* determine the ecological outcomes. If both invasion growth rates are positive (mutual invasibility), all three species coexist. If both invasion rates are negative, there is a priority effect in which both single resource-consumer equilibria are stable. If one invasion growth rate is positive and the other is negative, this suggests (as confirmed by numerical simulations) that the resource species with the positive invasion growth rate excludes the other resource species. In the supplementary material (Supplement A), we provide more details about this analysis and describe the invasion conditions for these outcomes or when the consumer only persists in the presence of both resource species.

As a step toward understanding the general conditions for coexistence and exclusion, we first examine two special cases corresponding to (i) resource competitive symmetry whereby *α*_11_ = *α*_12_ = *α*_21_ = *α*_22_ and (ii) pure apparent competition whereby *α*_12_ = *α*_21_ = 0. When there is competitive symmetry, the resource species with the larger value of *r_i_/A_i_* excludes the other resource species. The quantity *r_i_/A_i_* corresponds to the consumer density supported by resource species *i* when intraspecific competition is very weak i.e. *α_ii_* = 0. Hence, the resource species that supports the highest consumer equilibrium density (in the absence of self-limitation) excludes the other. This is an analog of the *P*\* rule (Holt 1977, Holt and Lawton 1993, Schreiber 2021). In this case, sexual dimorphism (as measured by *β*) influences outcomes only if there is an asymmetry in the maximal attack rates of the female and male (e.g., female masked boobies dive for prey more often than males; Weimerskirch et al. 2009). For example, if the female-preferred resource (*R_1_*) has the higher intrinsic rate of growth (*r*_1_ > *r*_2_) and the female consumer has the higher maximal attack rate of the two sexes, then the female-preferred resource species excludes the other resource species when sexual dimorphism is low (i.e. *A_1_=A_2_* when *β* = 0). However, as the average attack rate *A_1_* on the female-preferred resource increases with the magnitude *β* of the sexual dimorphism, sexual dimorphism reverses the outcome of apparent competition whenever *r*_1_/*a_F,max_* < *r*_2_/*a_M,max_* and the self-limitation in the female-preferred resource is weak.

Next, we consider the case of pure apparent competition in which there is no direct competition between the resources, i.e. *α*_12_ = *α*_21_ = 0. Then, coexistence occurs when

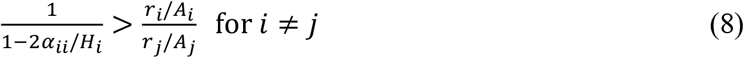

The right-hand side of this condition determines the outcome of apparent competition in the case of competitive symmetry i.e. the species with larger *r_i_/A_i_* wins. The left-hand increases to infinity as the sexual dimorphism increases such that *H_i_* approaches *2α_ii_*. This has two implications: (i) when *r*_1_/*A*_1_ and *r*_2_/*A*_2_ are equal, the resource species coexist at any level of sexual dimorphism, and (ii) when *r*_1_/*A*_1_ and *r*_2_/*A*_2_ are unequal, sufficiently strong sexual dimorphisms (that still allow the consumer to persist, per equation 5) will ensure coexistence of the two resource species by diluting the strength of apparent competition.

Finally, the general coexistence condition is

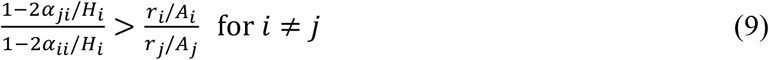

The effect of sexual dimorphism on the left-hand side term depends on the relative strengths of intra- and interspecific competition. When intraspecific competition is greater than interspecific competition (i.e. *α*_ii_ > *α_ji_*), the left-hand side term increases to infinity with increasing sexual dimorphism. In contrast, when interspecific competition between resources is greater than intraspecific competition (i.e. *α_ji_* > *α_ii_*), the left-hand side term of the general coexistence condition decreases to zero with increasing levels of sexual dimorphism. The effect of sexual dimorphism on the right-hand side term of the general coexistence criterion is as discussed for the case of competitive symmetry. Together these observations imply that if resource 1 is the better direct competitor i.e. *α*_21_ > *α*_11_and *α*_21_ < *α*_22_, then coexistence requires that resource 2 is the better apparent competitor i.e. *r*_2_/*A*_2_ > *r*_1_/*A*_1_. In which case, a sexual dimorphism can help consumer-mediated coexistence.

### Numerical results: Symmetric resource acquisition across the sexes

Our numerical results generally matched conclusion from our analytical solutions and are summarized in Table 1. The effects of consumer sexual dimorphism on resource population dynamics are illustrated in Figure 2. Sexual dimorphism results in increased resource density (Fig 2B vs. 2D; Fig 3), reduced consumer density (Figure 3), and can facilitate persistence of a competitively inferior resource species (Fig 2A vs. 2B; note that this contrast is determined by persistence of the consumer, which is possible under the conditions in 2B). When competition is purely apparent and resource acquisition by the consumer is symmetric across the sexes, increasing consumer sexual dimorphism increases the parameter space under which resources can coexist (Fig. 2C vs 2D), consistent with our analytical results (Fig. 4A). In particular, as sexual dimorphism approaches the extreme (each sex uses a different resource exclusively), competitively inferior resource species can coexist with a superior competitor that is more susceptible to apparent competition (Figure 4). This effect is in part due to the fact that consumer persistence depends on the density of both resources at this extreme.

**Figure 2.**
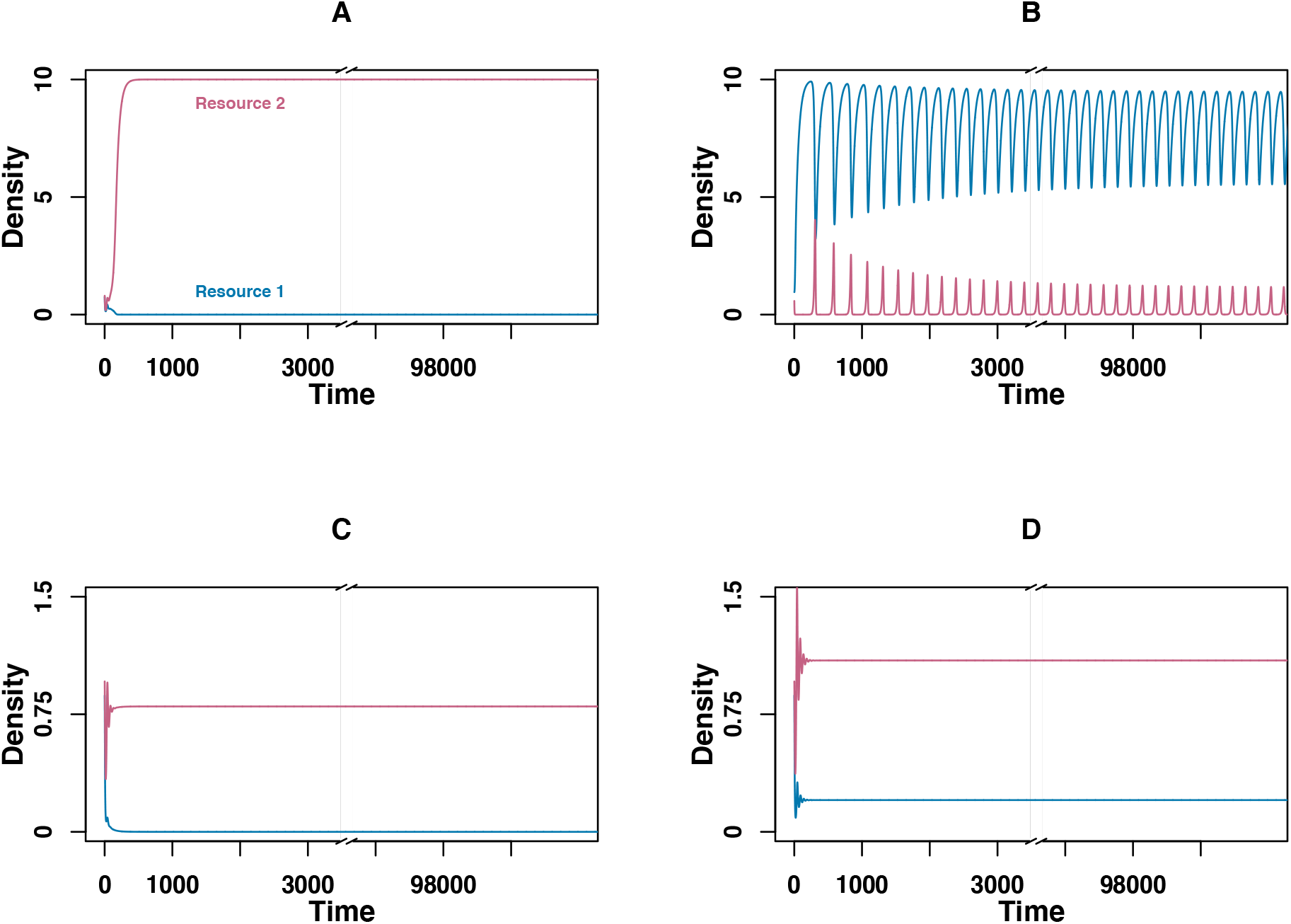
Coexistence mediated by sexual dimorphic consumers. In panel A, resource 2 (red) is a superior competitor to resource 1 (blue), leading to exclusion of resource 1 and the sexually-monomorphic consumer that cannot persist on a single resource species. However, adding consumer sex differences in total resource acquisition, under otherwise identical parameter values, leads to consumer persistence and coexistence of the resource species with cyclical dynamics (B). When competition is completely apparent (C, D), the resource with superior growth rate (red) excludes the inferior resource when consumers are sexually monomorphic (C). Under the same parameterization but with sexually dimorphic consumers (D), density of both resources is increased and they coexist. Parameter values: **A**, α_11_ = .1, α_12_ = .12, α_22_ = .1, α_21_ = .09, β = 1, *d_M_* = *d_F_* = 0.02, *b_M_* = *b_F_* = .1, *a_M, max_* = *a_F, max_* = 1, *r*_1_ = *r*_2_ = 1; **B** α_11_ = .1, α_12_ = .12, α_22_ = .1, α_21_ = .09, β = 1, *d_M_* = *d_F_* = 0.02, *b_M_* = *b_F_* = .1, *a_M, max_* = .8, *a_F, max_* = 1.2, *r*_1_ =1, *r*_2_ = 1; **C** α_11_ = .1, α_12_ = 0, α = .1, α_21_ = 0, β = 0, *d_M_* = *d_F_* = 0.02, *b_M_* = *b_F_*. 1, *a_M, max_* 1, *a_F, max_* 1, *r* = 1, *r*_2_ = 1.1; **D** α_11_ = .1, α_12_ = 0, α_22_ = .1, α_21_ = 0, β = 0.9, *d_M_ d_F_* 0.02, *b_M_ b_F_* .1, *a_M, max_* = 1, *a_F, max_* = 1, *r*_1_ = 1, *r*_2_ = 1.1;

**Table 1.** Summary of effects of sex differences in resource use. Arrows indicate effects on mean species density and the range of parameter space under which resource coexistence is observed.

| Density | Coexistence |
| --- | --- |
| Consumer ↓ | Apparent competition ↑ |
| Resource ↑ | Direct competition, equal resource growth rates, symmetric acquisition 0 |
|  | Direct competition, different resource growth rates, symmetric acquisition ↑ |
|  | Direct competition, asymmetric acquisition ↑↓ |

When resources compete directly with symmetric intraspecific competition, resource acquisition is symmetric across the sexes and intrinsic per-capita growth rates are equal between resources, consumer sexual dimorphism has little effect on invasion of a resource into a consumer-single-resource community at equilibrium (S1A-C, S4A). Yet even in the absence of effects on coexistence consumer sexual dimorphism has strong effects on equilibrium resource abundance, both under analytical solutions to a two species community and under simulations of three species communities (Fig 3). As predicted by our analytical conditions, when resource growth rates are unequal, sexual dimorphism changes expected regions of coexistence of both resources and the consumer, leading to invasion and persistence of competitively inferior resource that is a superior apparent competitor i.e. has a larger value of *r_i_/A_i_* (Figure 5B, Figure S4B, Figure S1).

**Figure 3.**
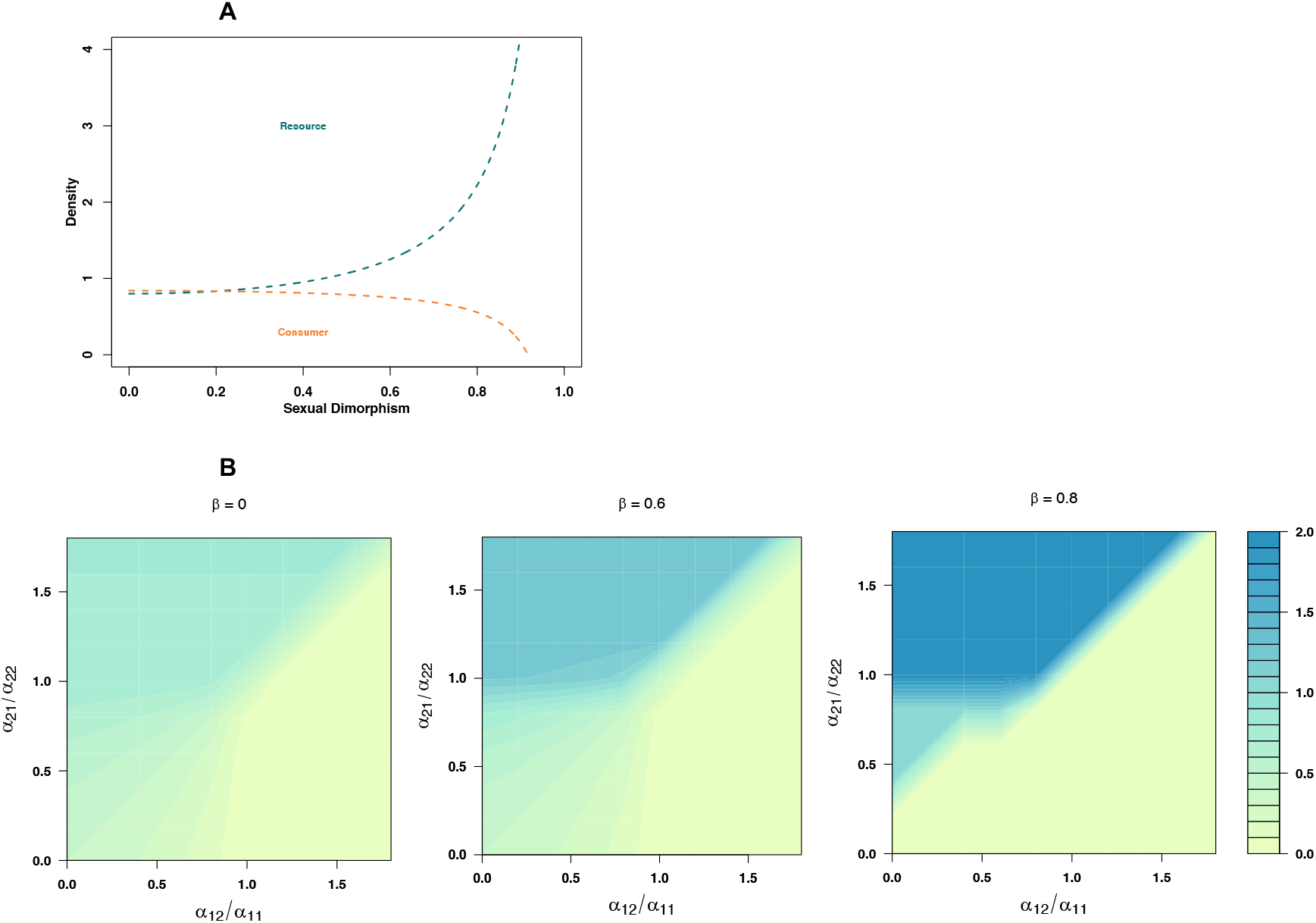
Consumer sexual dimorphism results in increased resource density and reduced consumer density in two and three species communities. Panel A: Equilibrium consumer female density decreases and resource 1 density increases with increased sexual dimorphism in a two species (consumer-1 resource) community (equation 6). Panel B: these results hold in simulations in three species communities, where resource 1 density is plotted against competition coefficients in both resource species under three levels of sexual dimorphism, increasing from left to right. Assuming symmetric total attack rates across the sexes, symmetric contribution of resource acquisition to consumer birth rates, and equal growth rates across resources.

### Numerical results: Asymmetric total resource acquisition

Introducing asymmetric total resource acquisition, represented as sex differences in the maximal attack rates *a_F,max_* and *a_M,max_*, has complex effects on both population dynamics and coexistence between competing resources. Introducing asymmetric attack rates can promote coexistence (as predicted by the analytical results) while simultaneously creating cyclical resource dynamics (Fig 2B). Under pure apparent competition and asymmetric attack rates, consumer sexual dimorphism has some similar effects to the case of symmetry, mediating coexistence of resources that differ in intrinsic per-capita growth rates (Fig 4B).

**Figure 4.**
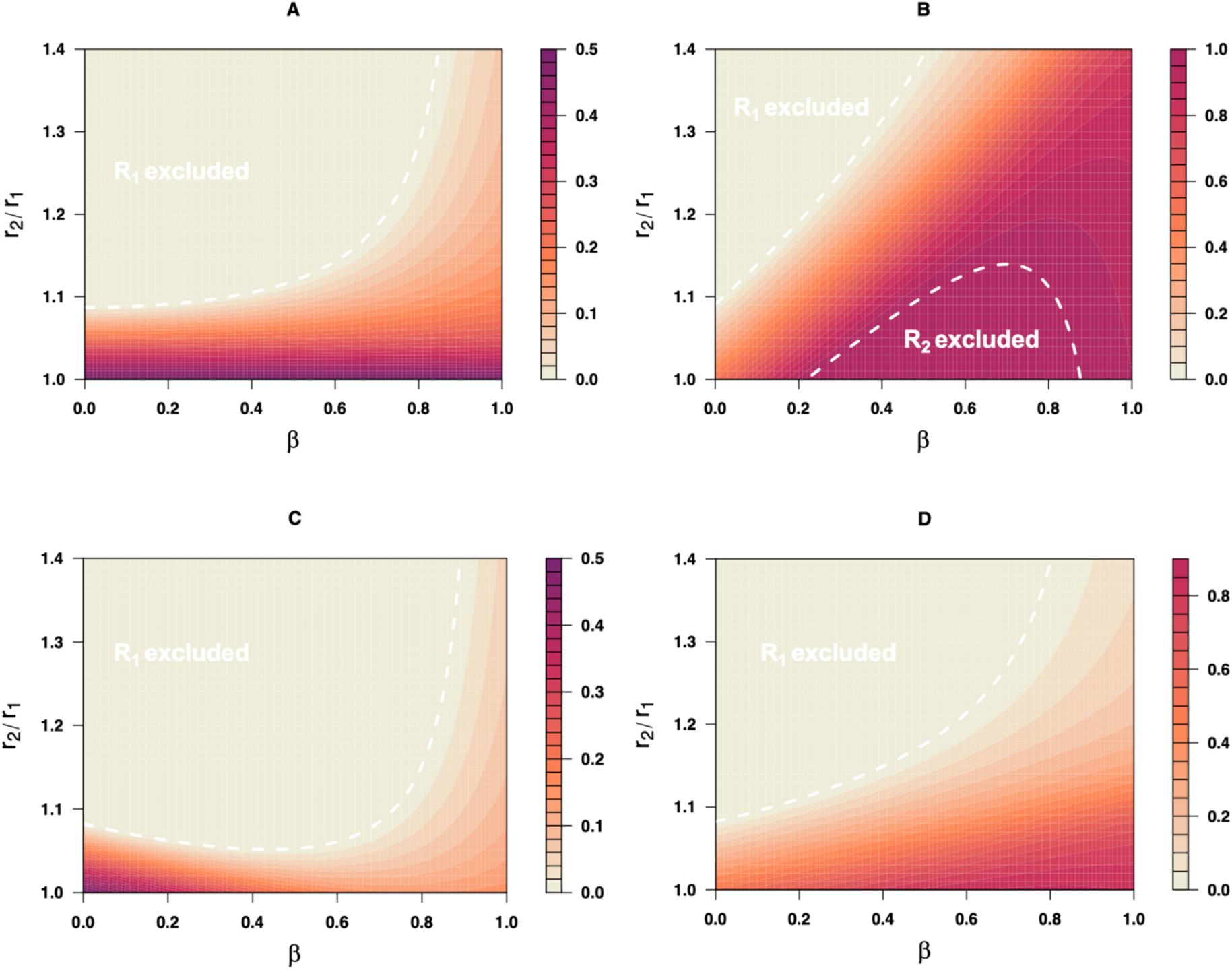
Sexual dimorphism in resource acquisition expands regions of coexistence between resources under apparent competition. Panels show relative density of resource 1 (*R_1_*/(*R_1_+R_2_*)) from numerical simulations with starting conditions of *R_1_ = R_2_ = 1* and *M = F = 1*. Panel A shows the outcomes under symmetric maximal attack rates across the sexes (*a_M, max_* = *a_F, max_* = 1), and equal death rates *d_M_* = *d_F_* = 0.02. Panel B shows the outcomes assuming sex differences in maximal attack rate (*a_M, max_* = .8, *a_F, max_* = 1.2). Panel C shows outcomes under identical conditions to A, but with a sex difference in death rate of approximately 11% (reduced male mortality, *d_M_* =0.018, *d_f_* =0.02). Panel D shows the opposite sex difference (reduced female mortality, *d_M_* =0.02, *d_f_*=0.018). White dashed lines indicate analytical solutions (equation 8) limited to the range of consumer coexistance 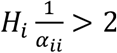, which corresponds to *β* < .96.

Under direct interspecific competition, sexual dimorphism in resource specialization has striking effects on resource persistence when the sexes differ in total resource acquisition, expanding regions of coexistence (Figs 5A, B) and in some cases leading to persistence of a competitively inferior resource (Figs 5C, D; Supplement B Figs S4B, C). Moreover, sex differences in total resource acquisition can reverse competitive outcomes when resources differ in their intrinsic per-capita growth rates and the degree of resource specialization is held constant (Figs 5C, D). The effect of sex differences in total resource acquisition on patterns of prey persistence can be as striking as the effects of resource specialization (Supplement B Figs S2, S1).

**Figure 5.**
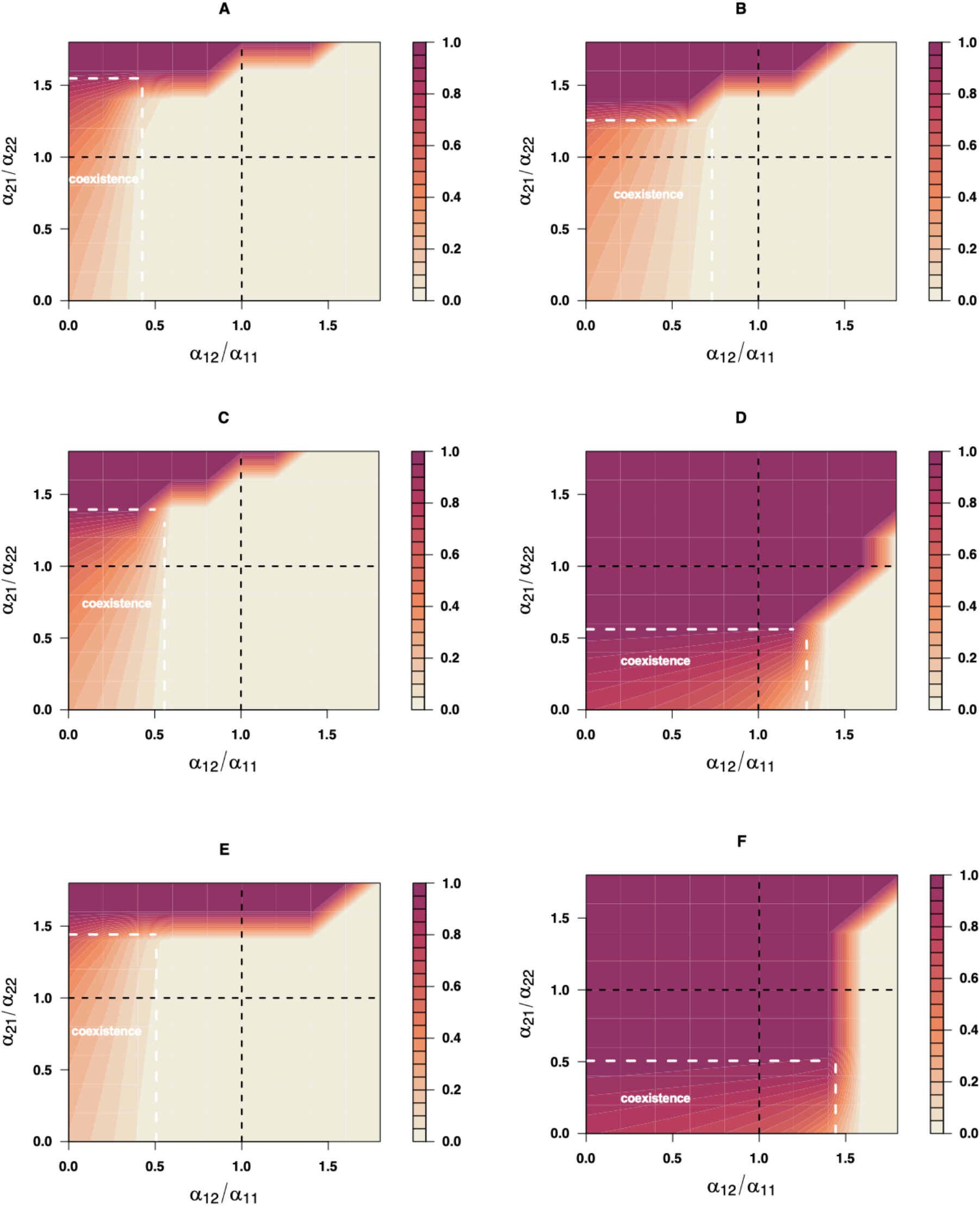
Sexual dimorphism alters regions of coexistence of resource species. Panels show relative density of resource 1 (*R_1_*/(*R_1_ + R_2_*)) from numerical simulations with starting conditions of *R_1_* = *R_2_* = 1 and *M=F* = 1. Panel A shows outcomes under sexual monomorphism (β = 0), unequal growth rates (*r_1_* = 2, *r_2_* = 2.1) across resources, and no sex differences in total attack rate (*a_M, max_* = *a_F, max_* = 1). Panel B shows the outcome under the same parameter values but with strong consumer sexual dimorphism (β = 0.7). Panels C and D show the effect of sexual niche divergence when the sexes also differ in total attack rates (total resource acquisition), with C showing the outcomes under unequal total attack rates (*a_M, max_* = .7, *a_F, max_* = 1.3), sexual monomorphism in prey preference (β = 0), and differential resource growth rates (*r_1_* = 1.6, *r_2_* = 1.8), and D showing the outcome under the same parameter values but with moderate sexual dimorphism in prey preference (β = 0.4). Panels E and F show the effects of sex differences in mortality; in Panel E, *d_M_* =0.018, *d_f_* =0.02. In Panel F, *d_M_* =0.02, *d_f_*=0.018, while β = 0.5 in both panels. Note that in the absence of sex differences in mortality coexistance would be restricted to the bottom left quadrant represented by black dashed lines, which demark equal inter and intraspecific competition coefficients. White dashed box demarks coexistence determined by analytical invasion criteria into a two-species community under equation 9.

### Asymmetric contribution of resource acquisition to birth rate

Introducing asymmetries in the contribution of sex-specific resource acquisition to birth rate (e.g., polygyny or polyandry, manifest as changes of the constant *b_M_*), had little qualitative effect. This similarity is illustrated in Supplement B figure S3 (compare to S1). Although changing these constants results in shifts of absolute regions of coexistence, the influence of consumer sex differences was similar. Lack of sensitivity in asymmetries in the contribution of resource acquisition to birth rates is consistent with our analytical results (Supplement A) that apply to all first-order homogenous mating functions including geometric and arithmetic contributions to reproduction.

## Discussion

Using a general model of consumer-resource dynamics we show that consumer sexual dimorphism has substantial consequences for community assembly. Competitive exclusion via apparent competition is expected and observed when males and females are monomorphic generalists and resources differ substantially in their intrinsic per-capita growth rates. However, when male and female consumers differ in their resource-specific attack rates, resource species that differ substantially in their intrinsic per-capita growth rates can coexist. Similar effects of consumer sexual dimorphism are observed when resources compete directly, with sexual dimorphism in some cases permitting coexistence or persistence of a competitively inferior resource (summarized in Table 1). However, for both direct and apparent competition, consumer sexual dimorphism can also lead to competitive exclusion between resources that would typically be expected to coexist. Thus, consumer sex differences result in fundamental changes in the types of competing resources that can establish during community assembly. These results also support conclusions from other food web models suggesting trophic position may impact the observed ecological effects of sexual reproduction (Kawatsu 2018). Moreover, equilibrium resource abundances and temporal dynamics are altered by consumer sexual dimorphism even when long-term ecological outcomes are unaffected. Our results in many ways echo recent work demonstrating that ontogenetic differences in resource acquisition in a species can have similar complex consequences for community assembly (de Roos 2020). However, a key difference is that in sexually-reproducing species we expect to observe these types of effects whenever consumer mating propensity depends, in part, on resource acquisition, a biological reality for many sexually-reproducing consumers (e.g. widespread evidence of developmental thresholds for transitions to reproduction, reveiwed in Wilbur and Collins 1973, Day and Rowe 2002, and resource-dependent sexual displays, Bonduriansky 2007), and whenever male abundance matters for population growth rates.

Although our results suggest that the effects of consumer sexual dimorphism on community assembly can be complex, our analysis reveals some key predictions from our model. First, consumer sex differences in resource-specific attack rates result in an increase in the equilibrium density of each resource and a decrease in the abundance of the consumer species. Intuitively, sexual dimorphism can frequently lead to slightly suboptimal resource use of the species due to a mismatch between demand (restricted by a 50:50 sex ratio) and resource availability which can be more variable and dynamic. Second, consumer sexual dimorphism can promote coexistence when the more aggressively-feeding sex (higher total attack rates) specializes on and suppresses what would otherwise be the competitively superior resource. This later situation may be commonplace if sex-specific natural selection favors specialization, by the sex with higher resource requirements, on the most abundant resource. Finally, and related to the previous observation, the effects of consumer sexual dimorphism are most pronounced when resources either differ in intrinsic per-capita growth rates, or when consumer sexes differ in their total resource acquisition.

Although the assumption of complete female demographic dominance is common in theoretical models in ecology and evolutionary biology, there is little empirical support for this demographic extreme. Our simulation models instead focused on a harmonic mean birth function, which is biologically realistic in that it captures severely reduced birth rates when any one sex is rare, which is likely to be the case even if sperm limitation is unimportant at sex ratios near 1:1. Empirically evaluating alternative birth functions in real systems is difficult, requiring extreme variation in adult sex ratios rarely seen in many species. However, some empirical data exists and supports the harmonic mean as the best-fitting description of sexual birth rates (Miller and Inouye 2011a). Nonetheless, our analytical results (see Supplement A for details) that our conclusions apply to any birth function that is first order homogenous i.e. *B*(*aMI_M_, aFI_F_*) = *aB*(*MI_M_, FI_F_*) for any positive constant *a*>0. These functions include all functions typically considered candidate birth functions by demographers (Caswell and Weeks 1986, Miller and Inouye 2011a). Our conclusions do depend on the assumption that male abundance makes some contribution to birth rate.

Although even moderate sex differences in attack rates can change the conditions under which resources coexist, the strongest effects are observed when the consumer population approaches complete sexual dimorphism in resource specific attack rates. Cases of such extreme sex differences in diet preference exist (Temeles et al. 2000), although whether they occur with any regularity is unclear. Diet divergence between the sexes, and sexual dimorphisms in trophic morphology, are common (Shine 1989), although overlap in male and female diets can be substantial even in the presence of ecological sex differences (e.g., Stamps et al. 1997, De Lisle and Rowe 2015a). Yet overlap in diet content may be expected even in the case of large differences in attack rates, as diet content is the product of both attack rate and resource abundance. Further, males and females may fail to diverge in diet preference even when optimum diet nutritional content differs across the sexes (Reddiex et al. 2013). These challenges to understanding sex-specific resource acquisition highlight the need for further empirical studies disentangling expressed diet preference, male and female nutritional optima, and the evolution of ecological sexual dimorphism.

We have assumed a Fisherian sex ratio, where the primary sex ratio is maintained at a stable 1:1 ratio (Fisher 1930), although our model allows for deviations from a 1:1 operational sex ratio via sex-specific intrinsic mortality. In fact, our results show that even slight sex differences in mortality rates can have dramatic consequences for community assembly under both pure apparent and direct competition between resource species. Sexspecific mortality is commonplace in natural populations and can occur for a variety of reasons, such as sex-specific costs of reproduction, predation, or sexual conflict. In our model, sex-specific mortality, as well as variation in consumer mating system (see Supplemental material) had strong but predictable consequences for resource abundance, leading to increases in the abundance of the resource favored by the sex with higher intrinsic mortality.

Our analysis also assumes that the male and female trophic traits are constant. In nature, however, the degree and even direction of sexual selection can vary dramatically among closely related populations (e.g., Reimchen et al. 2016). Dimorphism clearly evolves rapidly in response to spatially varying sexual selection and resource availability. Future extensions to our strictly ecological model could add in eco-evolutionary dynamics of sexual dimorphism.

Male and female densities were reduced with increasing ecological sexual dimorphism, because consumer births are limited by resource acquisition in both sexes in our model. In the extreme case, where the consumer cannot persist on a single resource (for example, under complete sexual dimorphism), coexistence of the consumer depends on stable coexistence of the resource species (Supplement A). Our simulations suggest robust persistence of extremely dimorphic consumers in two-consumer communities, albeit at reduced density. Both of these results – reduced consumer density and dependence on the presence of both resource species in the community – suggest that sexually-dimorphic populations may face a higher risk of extinction due to demographic stochasticity. However, it is difficult from our purely deterministic ecological model to fully interpret the consequences of sexual dimorphism on extinction probability. Evolution of sexual dimorphism from a monomorphic ancestor that specializes on a single resource could lead to increased population mean fitness, by increasing the total resource pool available across both sexes (Rand 1952, Selander 1966, Slatkin 1984, Li and Kokko 2021); such a process represents a form of within-species, between-sex ecological character displacement, and may be particularly likely to occur when the sexes interact in small demes (Li and Kokko 2021). More generally, evolution of sexual dimorphism is expected to be critical to population mean fitness whenever optima differ for males and females (Lande 1980), and empirical data suggest sexual dimorphism can be associated with reduced extinction probability at both the macroevolutionary scale and in extant populations (De Lisle and Rowe 2015b). Nonetheless, our results suggest a full understanding of sexual dimorphism’s role in population persistence could require integrating theory and data from population/community ecology and evolutionary genetics. This conclusion is complemented by recent theory (de Vries and Caswell 2019) indicating sexual dimorphism in demographic parameters can have important consequences for maintaining genetic diversity.

Our model generally predicts that ecological sexual dimorphism may in some cases promote, and in other cases reduce, diversity at lower trophic levels during community assembly. Testing this prediction with correlative data (reviewed in Tsuji and Fukami 2020) would be possible but challenging. An alternative and non-exclusive hypothesis, that dimorphic predators are more likely to establish in communities with diverse prey assemblages, would also generate patterns consistent with our results. Alternatively, experiments that manipulate the expressed dimorphism of predators and track community dynamics at lower trophic levels are tractable in some systems. Similar experiments have been performed (Fryxell et al. 2015, Start and De Lisle 2018), in which effects of predator sex ratio manipulation on prey communities are assessed in mesocosm designs. Although the results of these experiments do suggest sex differences can have important community consequences, no studies (to our knowledge) have compared communities in which the magnitude of morphological sexual dimorphism is manipulated under a stable sex ratio. Such designs are possible when distributions of male and female phenotypes exhibit substantial variation, and would represent an ideal empirical test of the theoretical results presented here.

Ecological sex differences are commonplace, although the details of their evolutionary drivers and ecological consequences are unclear. Emerging theory and data indicate ecological sex differences may have important consequences for the evolutionary genetics of adaptation (Zajitschek and Connallon 2017), the dynamics of diversification (Bolnick and Doebeli 2003, De Lisle and Rowe 2015b) and community assembly (Fryxell et al. 2015, Pincheira-Donoso et al. 2018, Start and De Lisle 2018). Our results add to this body of work, indicating that sexual dimorphism can have substantial effects on the structure, abundance, and dynamics of ecological communities, including changing conditions for coexistence between competing resource species.

## Acknowledgements

We thank Tim Connallon and Miguel Gomez for comments on an early draft of this manuscript, as well as Gonzalo Hernando for assistance in the early stages of the project. This work was supported by funds from the University of Connecticut to DIB, an establishment grant from the Swedish Research Council to S.P.D. (2019-03706), and an U.S. National Science Foundation Grant DMS176803 to SJS.

