## Supplement A for "Complex community-wide consequences of consumer sexual dimorphism"

**Supplement A: Invasion analysis** for "Complex community-wide consequences of consumer sexual dimorphism" by Stephen P. De Lisle, Sebastian J. Schreiber, and Daniel I. Bolnick

Let $R_i$ for $i = 1, 2$ be the density of resource species $i$, and $F, M$ be the female and male consumer densities, respectively. In the absence of the consumer, the resources exhibit classical Lotka-Volterra competitive dynamics with intrinsic rates of growth $r_i$ and competition coefficients $\alpha_{ij}$. The attack rates of the female and male consumers on resource $i$ equal $a_{Fi}$ and $a_{Mi}$, respectively. The per-capita death rates of the females and males are $d_F, d_M$, respectively. As explained in the main text, the consumer birth rate depends on the captured resources, specifically the functions $I_M = \sum_i b_{Mi}a_{Mi}R_i$ and $I_F = \sum_i b_{Fi}a_{Fi}R_i$. We assume the birth rate equals $B(MI_M, FI_F)$ where $B(x, y)$ is a non-negative, first-order homogeneous function i.e. $B(ax, ay) = aB(x, y)$ for any $a \geq 0$. Examples of functions satisfying this assumption include the harmonic mean mating function $B(x, y) = \frac{1}{\frac{1}{2}x+\frac{1}{2}y} = \frac{2xy}{x+y}$, the linear mating function $B(x, y) = \alpha x + \beta y$ with $\alpha, \beta \geq 0$, and the geometric mean mating function$B(x, y) = \sqrt{xy}$. The results in the main text focus on the case of the harmonic mean mating function.

Under these assumption, the model for the consumer-resource dynamics is

$$\begin{aligned}\frac{dR_i}{dt} =& r_iR_i(1 - \alpha_{i1}R_1 - \alpha_{i2}R_2) - a_{Mi}MR_i - a_{Fi}FR_i \qquad i = 1, 2\\ \frac{dF}{dt} =& \frac{1}{2}B(MI_M, FI_F) - d_FF\\ \frac{dM}{dt} =& \frac{1}{2}B(MI_M, FI_F) - d_MM\end{aligned} \tag{A1}$$

To derive the conditions for coexistence, exclusion, and bistability, we perform an invasion analysis consistent with the theory of permanence (Schreiber 2000, Patel & Schreiber 2018). The invasion analysis focuses on two cases: (i) the consumer can persist on each resource species, and (ii) the consumer can persist on neither resource species. In case (i), we solve for the consumer-resource $i$ equilibria and determine the per-capita growth rate of the missing resource species $j \neq i$. These per-capita growth rates, known as invasion growth rates, determine whether or not the resource species can increase when rare or not. In case (ii), coexistence (in the sense of permanence) is only possible if the two resource species coexist in the absence of the consumer and the consumer species has a positive per-capita growth rate at this two species coexistence equilibrium.

**Case (i):** Assume the consumer and resource $i$ subsystem persists. The coexistence equilibrium of this subsystem satisfies

$$\frac{1}{2}B(MI_M, FI_F) - d_F F = 0 = \frac{1}{2}B(MI_M, FI_F) - d_M M$$

which implies

$$M = \frac{d_F}{d_M}F$$

at this equilibrium. Using this relationship and the $\frac{dF}{dt} = 0$ condition, we get

$$\frac{1}{2}B\left(\frac{d_F}{d_M}FI_M, FI_F\right) = d_F F$$

at the coexistence equilibrium. Moreover, as $I_M = b_{Mi}a_{Mi}R_i$ and $I_F = b_{Fi}a_{Fi}R_i$ in the two species system. Therefore,

$$\frac{1}{2}B\left(\frac{d_F}{d_M}Fb_{Mi}a_{Mi}R_i, Fb_{Fi}a_{Fi}R_i\right) = d_F F$$

at the coexistence equilibrium. Homogeneity of $B$ implies

$$\frac{FR_i}{2}B\left(\frac{d_F}{d_M}b_{Mi}a_{Mi}, b_{Fi}a_{Fi}\right) = d_F F.$$

Dividing by $F$ and solving yields the equilibrium value of $R_i$

$$R_i^* = \frac{2d_F}{B\left(\frac{d_F}{d_M}b_{Mi}a_{Mi}, b_{Fi}a_{Fi}\right)} = \frac{2}{B\left(\frac{b_{Mi}a_{Mi}}{d_M}, \frac{b_{Fi}a_{Fi}}{d_F}\right)}$$

where the second equality follows from dividing numerator and denominator by $d_F$ and using homogeneity of $B$.

To determine the values of $F$ and $M$ at the coexistence equilibrium, we use the equilibrium condition for the resource equation

$$0 = \frac{1}{R_i}\frac{dR_i}{dt} = r_i(1 - \alpha_{ii}R_i^*) - a_{Mi}M - a_{Fi}F$$

As $M = \frac{d_F}{d_M}F$ at the coexistence equilibrium, we can solve for $F$ and $M$ to get

$$\begin{aligned} F^* =& \frac{r_i(1 - \alpha_{ii}R_i^*)}{\frac{d_F}{d_M}a_{Mi} + a_{Fi}} \\ M^* =& \frac{d_F}{d_M}F^* \end{aligned}$$

as presented in the main text.

The per-capita growth rate, call it $I_j$, of other resource species $j \neq i$ at this two species coexistence equilibrium is

$$I_j = r_j(1 - \alpha_{ij}R_i^*) - a_{Mj}M^* - a_{Fj}F^*.$$

Substituting in the equilibrium expressions and simplifying yields

$$I_j = r_j(1 - \alpha_{ij}R_i^*) - \frac{a_{Mj}\frac{d_F}{d_M} + a_{Fj}}{a_{Mi}\frac{d_F}{d_M} + a_{Fi}}r_i(1 - \alpha_{ii}R_i^*).$$

Thus, $I_j$ is positive if and only if

$$\frac{r_j(1-\alpha_{ij}R_i^*)}{r_i(1-\alpha_{ii}R_i^*)} > \frac{a_{Mj}\frac{d_F}{d_M}+a_{Fj}}{a_{Mi}\frac{d_F}{d_M}+a_{Fi}}$$

as presented in the main text.

**Case (ii)** Consider the case where the consumer can not persist on either resource alone i.e. $B\left(\frac{b_{Mi}a_{Mi}}{d_M}, \frac{b_{Fi}a_{Fi}}{d_F}\right)/\alpha_{ii} < 2$ for $i = 1, 2$. In this case, coexistence (in the sense of permanence) is only possible if the resource species (in the absence of the consumer) are able to coexist, say at the equilibrium $\hat{R}_1 > 0, \hat{R}_2 > 0$. To determine whether the consumer has a positive per-capita growth rate at this equilibrium, we make a change of variables by defining $x = F/(M + F)$ as the sex ratio and $C = M + F$ as the total consumer density. In this coordinate system, the consumer dynamics are

$$\begin{aligned}\frac{dC}{dt} =& C(B((1-x)I_M, xI_F) - d_F x - d_M(1-x))\\ \frac{dx}{dt} =& (0.5-x)B((1-x)I_M, xI_F)) + x(1-x)(d_M - d_F).\end{aligned}$$

Importantly, the per-capita growth rate of the consumer is

$$B((1-x)I_M, xI_F) - d_F x - d_M(1-x).$$

When the resources are at their equilibrium densities $\hat{R}_i$, we have $I_F^* = \sum_i b_{Fi}a_{Fi}\hat{R}_i$, $I_M^* = \sum_i b_{Mi}a_{Mi}\hat{R}_i$ and the sex ratio dynamics become

$$\frac{dx}{dt} = (0.5-x)B((1-x)I_M^*, xI_F^*)) + x(1-x)(d_M - d_F).$$

Numerical simulations suggest for the mating functions considered here, these sex ratio dynamics converge to a unique equilibrium $x^*$ in which case the per-capita growth rate of the consumer at the two species coexistence equilibrium equals

$$I_C = B((1-x^*)I_M^*, x^* I_F^*) - d_F x^* - d_M(1-x^*).$$

Coexistence requires that $I_C > 0$. In general, there is no simple analytic formula for $x^*$. However, in the special case that $d_M = d_F$, we get $x^* = 0.5$ and

$$I_C = \frac{1}{2} B(I_M^*, I_F^*) - d_F.$$
