## Supplement B for "Complex community-wide consequences of consumer sexual dimorphism"

**Supplement B: Supplemental figures and tables, and Development of Harem size birth function** for "Complex community-wide consequences of con- sumer sexual dimorphism" by Stephen P. De Lisle, Sebastian J. Schreiber, and Daniel I. Bolnick

### Table S1a. Parameter values for simulation runs presented in text

| Parameter | Description | Values |
|---|---|---|
| $\beta$ | Sexual dimorphism in prey preference | 0 - 1 |
| $r_1$ | $R_1$ intrinsic per-capita growth rate | 1 - 2 |
| $r_2$ | $R_2$ intrinsic per-capita growth rate | 1-2.1 |
| $\alpha_{11}, \alpha_{22}$ | intraspecific competition coefficients | 0.1 |
| $\alpha_{12}, \alpha_{21}$ | interspecific competition coefficients | 0 - 0.18 |
| $d_M\, d_F$ | consumer per-capita death rates | 0.001, 0.02 |
| $b_M$, $b_F$ | male & female contributions to consumer birth rate | 0.05-0.15 |
| $a_{M,\, max}$ | maximal attack rate of male consumers | 0.7 – 1 |
| $a_{F,\, max}$ | maximal attack rate of female consumers | 1 – 1.3 |
| $h$ | *Harem size* | 0.5, 1, 2 |

**Table S1b. Rescaled parameter values explored (results not shown, qualitative conclusions identical)**

| Parameter | Description | Values |
|---|---|---|
| $\beta$ | Sexual dimorphism in prey preference | 1, 0.999, .99, 0.9, 0.7, 0.5, 0.2, 0 |
| $r_1$ | $R_1$ intrinsic per-capita growth rate | 1, 1.5 |
| $r_2$ | $R_2$ intrinsic per-capita growth rate | 1, 1.0001, 1.001, 1.01, 1.1, 1.2, 1.4, 1.6 |
| $\alpha_{11}, \alpha_{22}$ | intraspecific competition coefficients | 0.0001, 0.00001 |
| $\alpha_{12}, \alpha_{21}$ | interspecific competition coefficients | 0, .00001, .00002, .00004, 0.00008, .0001, .00011, .00012, .00016, .00018, .000001, .000002, .000004, 0.000008, .00001, .000011, .000012, .000016, .000018 |
| $d_M\, d_F$ | consumer per-capita death rates | 0.002, 0.02 |
| $b_M$ | male contribution to consumer birth rate | 1, 0.8, 0.5, 0.3, 0.1, .0002 |
| $b_F$ | female contribution to consumer birth rate | 1, .0002 |
| $a_{M,\,max}$ | maximal attack rate of male consumers | 0.008, 0.007, 0.006, 0.004, 0.001 |
| $a_{F,\,max}$ | maximal attack rate of female consumers | 0.008, 0.009, 0.01, 0.012, 0.015 |

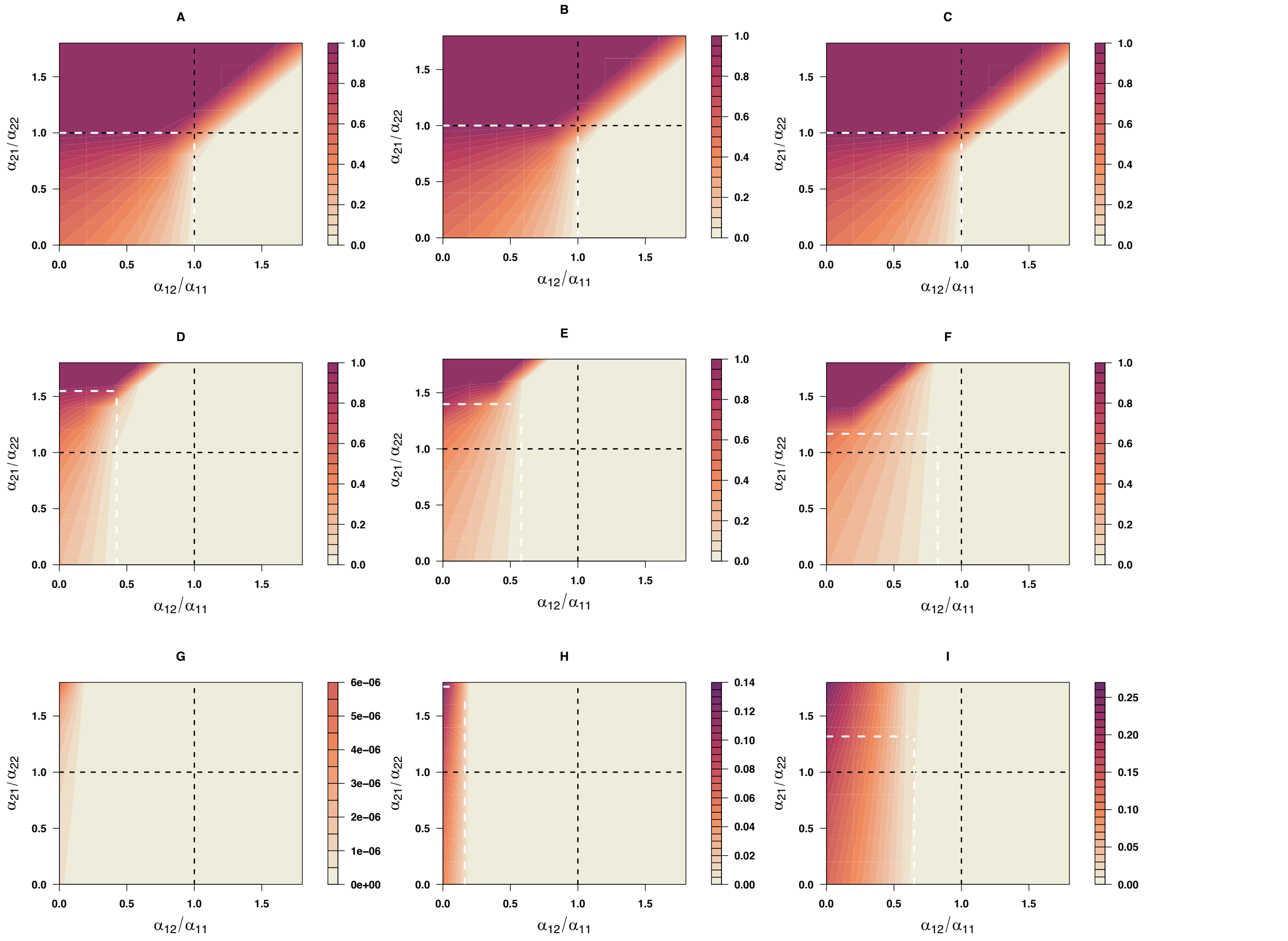


**Figure S1. Regions of persistence of the consumer/resource system assuming symmetric total resource acquisition and contribution of resource acquisition to birth rates.** Panels show relative density of resource 1 ($R_1$/( $R_1 + R_2$) from numerical simulations with starting conditions of $R_1 = R_2 = 1$ and $M = F = 1$. Parameter values: A, D, G $\beta = 0$, B, E H, $\beta = .5$, C, F I, $\beta = .8$, with A, B,C having $r_1 = r_2 =1$, D, E, F having $r_1 = 1$, $r_2 = 1.05$ and G, H, I having $r_1 = 1$, $r_2 = 1.1$. All other values were as listed in table S1A. Black dashed lines demark equal inter and intraspecific competition coefficients. White dashed lines demark analytical invasion criteria into a two-species community under equation 9.

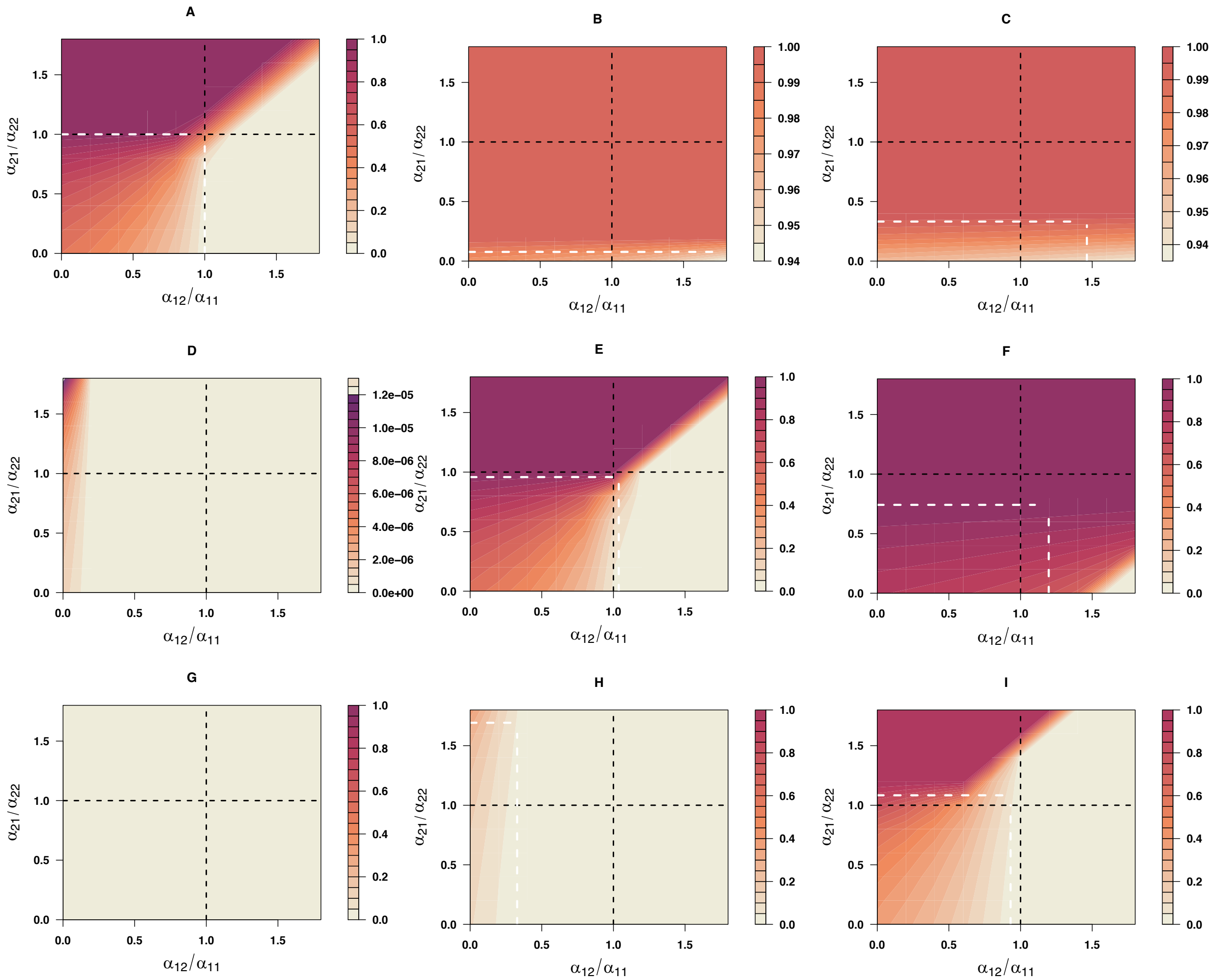


**Figure S2 Regions of persistence of the consumer/resource system assuming asymmetric total resource acquisition and symmetric contribution of resource acquisition to birth rates** Panels show relative density of resource 1 ($R_1/(R_1 + R_2)$) from numerical simulations with starting conditions of $R_1 = R_2 = 1$ and $M = F = 1$. In all panels, $a_{M,\,max}$ = .9, $a_{F,\,max}$ = 1.1. Parameter values: A, D, G β = 0, B, E H, β = .5, C, F I, β = .8, with A, B,C having $r_1 = r_2$ =1, D, E, F having $r_1$ = 1, $r_2$ = 1.05 and G, H, I having $r_1$ = 1, $r_2$ = 1.1. All other values were as listed in table S1A. Black dashed lines demark equal inter and intraspecific competition coefficients. White dashed lines demark analytical invasion criteria into a two-species community under equation 9.

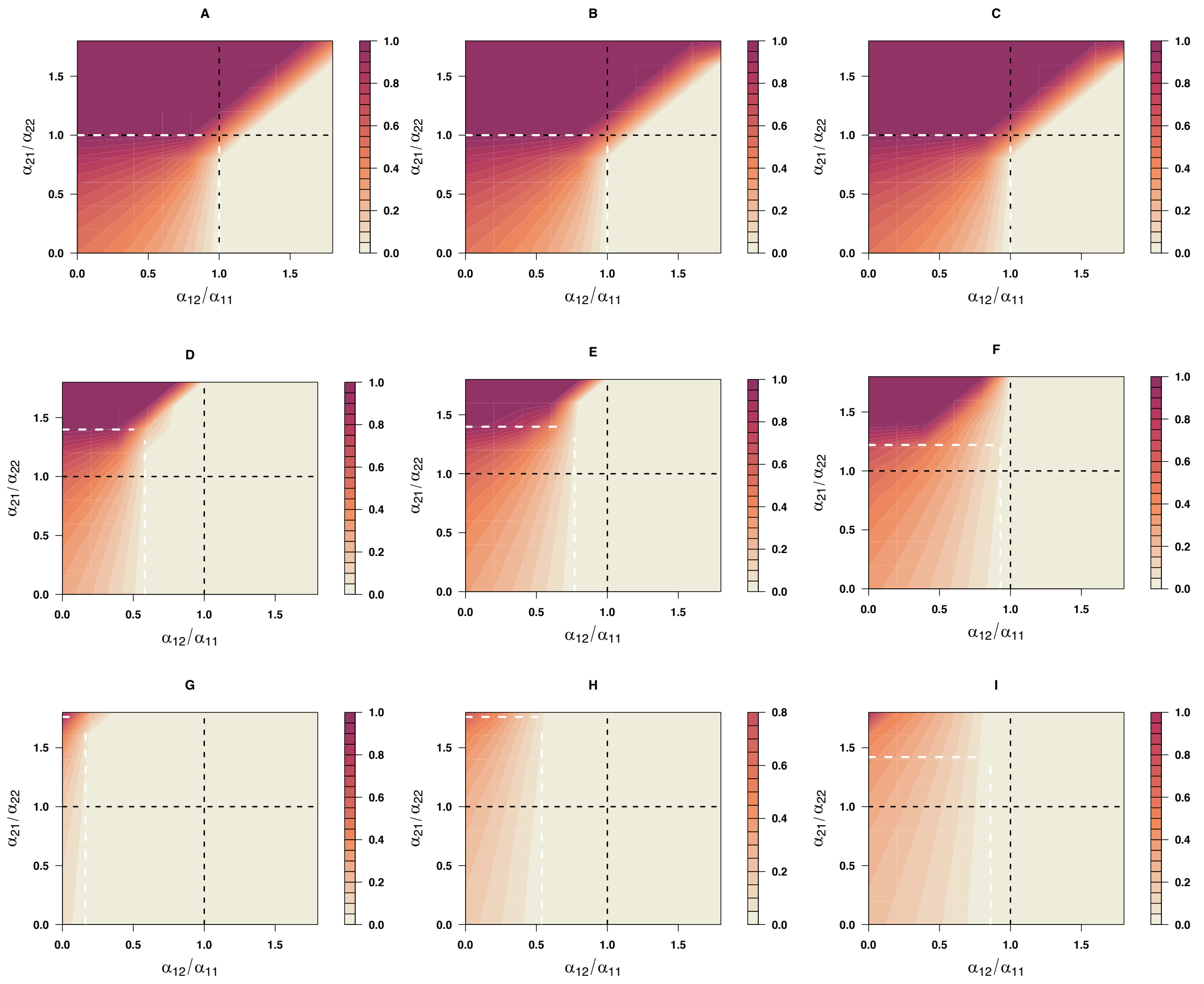


**Figure S3 Regions of persistence of the consumer/resource system assuming symmetric total resource acquisition and asymmetric contribution of resource acquisition to birth rates.** Panels show relative density of resource 1 ($R_1$/( $R_1$+ $R_2$) from numerical simulations with starting conditions of $R_1 = R_2 = 1$ and $M = F = 1$. In all panels, $b_M = .05$, $b_F = .15$, $a_{M,\ max} = a_{F,\ max} = 1$. Parameter values: A, D, G β = 0, B, E H, β = .5, C, F I, β = .8, with A, B,C having $r_1 = r_2 = 1$, D, E, F having $r_1 = 1$, $r_2 = 1.05$ and G, H, I having $r_1 = 1$, $r_2 = 1.1$. All other values were as listed in table S1A. Black dashed lines demark equal inter and intraspecific competition coefficients. White dashed lines demark analytical invasion criteria into a two-species community under equation 9.

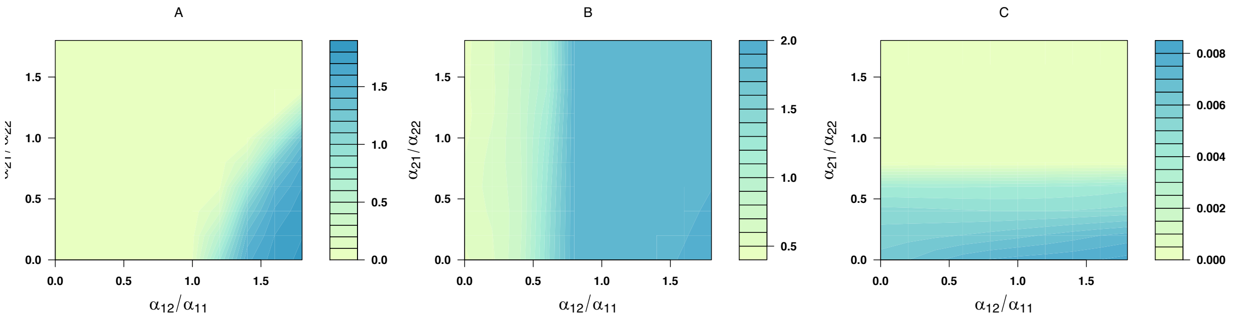


**Figure S4. Consumer sexual dimorphism can expand regions of resource invasion success into an equilibrium community.** Panels show density of resource 2 following invasion into a single resource-consumer community at equilibrium (see text) when consumer sexual dimorphism approaches completeness ($\beta = 0.99$). Panel A shows the case of equal total attack rates across the sexes ($a_{M,\,max} = a_{F,\,max} = 1$) and equal resource growth rates ($r_1 = r_2 = 2$). Panel B shows the outcome under the same values as A but assuming unequal resource growth rates ($r_1 = 2$, $r_2 = 2.1$). Panel C shows outcomes under the same parameter values as B but assuming sex differences in total attack rates ($a_{M,\,max} = .95$, $a_{F,\,max} = 1.05$). Other parameter values for all panels: $d_M = d_F = 0.001$, $b_M = b_F = 0.1$.

**Harem size**

Following (Caswell and Weeks 1986, Lindstöm and Kokko 1998), we can modify our birth function to include a harem size parameter *h* (where *h* > 1 corresponds to polygyny),

$$B(M,F,R_i,R_j) = 2\frac{MI_M * FI_F}{MI_M + FI_F/h}$$

$$=2\frac{MA_M*FI_F}{MA_M+FA_Fb_F/b_Mh}$$

where $A_M = I_M/b_M$. Setting $b_M h \rightarrow b_M$ yields

$$2\frac{MA_M * FI_F}{MA_M + FA_Fb_F/b_M}$$

$$= 2\frac{MI_M * FI_F}{MI_M + FI_F}$$

And thus including harem size *h* can be mathematically equivalent to modelling sex specific *b* (i.e., $b_M h$). Variation in *h,* equivalently sex specific *b*, couples births to on sex more than the other; when *h* > 1 birth rates are maximized under a female-biased sex ratio, and the opposite is true for *h* < 1 (Caswell and Weeks 1986). Variatin in *h* can shift regions of resource persistance in a corresponding manner, in the direction of the resource favored by females or males, respectively (Figure S5), similar to the effects of sex-speific *b* (Figure S6).

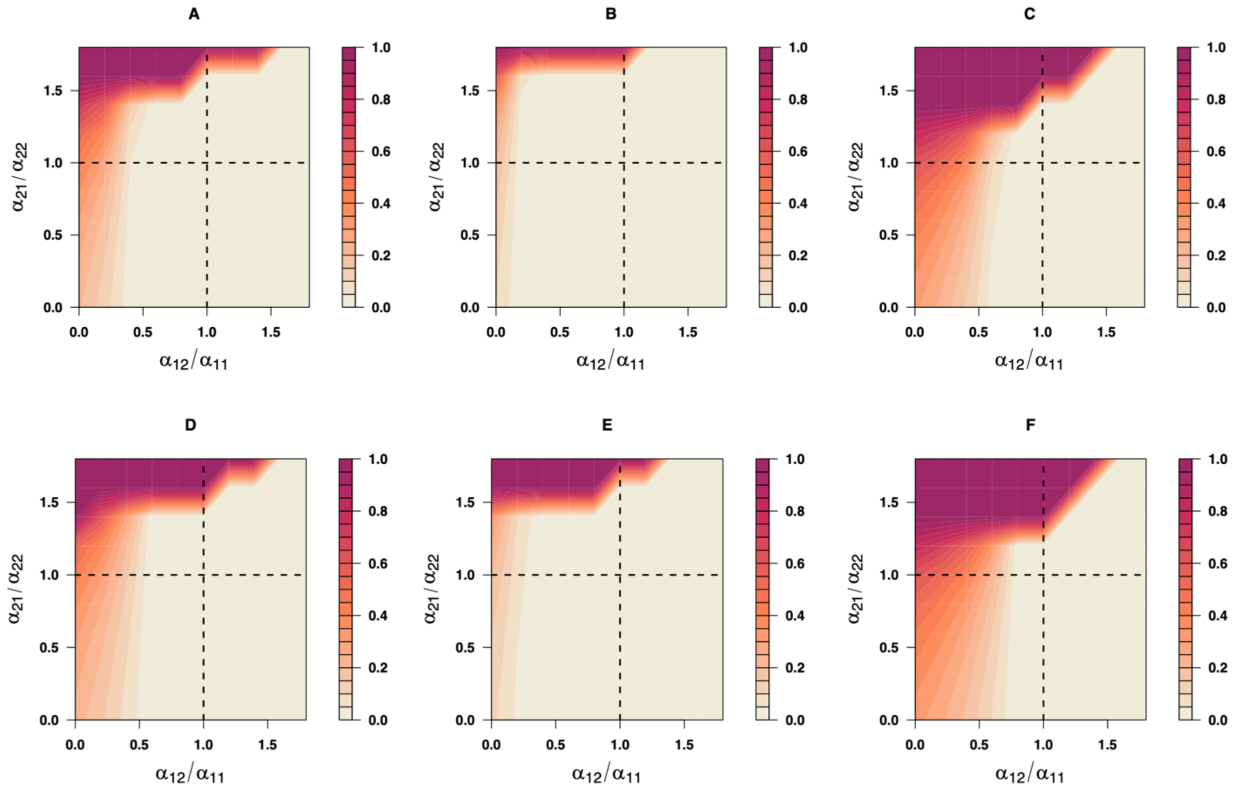


**Figure S5. Regions of persistence of the consumer/resource system under varying harem size h.** Panels show relative density of resource 1 ($R_1$/( $R_1 + R_2$) from numerical simulations with starting conditions of $R_1 = R_2 = 1$ and $M = F = 1$. On the left (A and D) $h$ = 1, middle (B and E) $h$ = 2 (females more important for birth rates), and right (C and F) $h$ = 0.5 (males more important for birth rates. Other parameter values: A, B, C β = 0.1; D, E F, β = .5, with all having $r_1 = 2$, $r_2$ =2.1, $b_M = b_F = 0.1$, $d_M = d_F = 0.02$. All other values were as listed in table S1A. Black dashed lines demark equal inter and intraspecific competition coefficients.

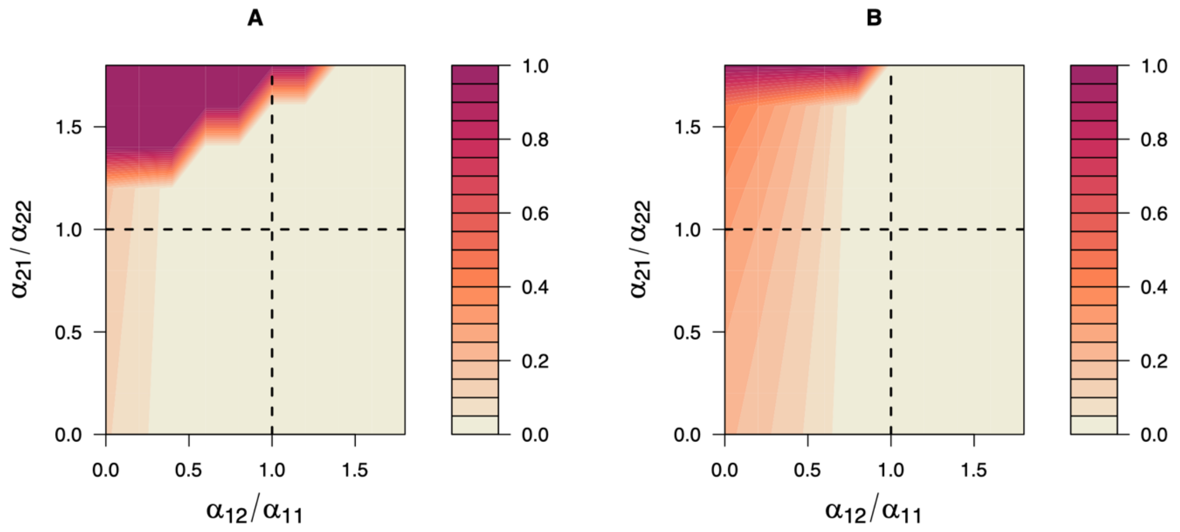


**Figure S6. Sex differences in b shift regions of persistence of the consumer/resource system.** Panels show relative density of resource 1 ($R_1/(R_1 + R_2)$) from numerical simulations with starting conditions of $R_1 = R_2 = 1$ and $M = F = 1$. In panel A, $b_M = 0.3$, $b_F = 0.1$; Panel B, $b_M = 0.1$, $b_F = 0.3$. Other parameters (same in both panels): $\beta = .8$, with all having $r_1 = 2$, $r_2 = 2.1$, $d_M = d_F = 0.02$.
